# Computational prediction of gamma-oryzanol as potential agonist of human peroxisome proliferator-activated receptor gamma (ppar-γ)

**DOI:** 10.1101/2023.06.10.542647

**Authors:** Prasanta Kumar Biswal, Swagatika Behera, Snehamayee Mohapatra, Durga Madhab Kar, Luna Samanta, Pabitra Mohan Behera, Rajat Kumar Kar

**Affiliations:** Department of Pharmacology, Dadhichi College of Pharmacy, Cuttack, Odisha, Pin-754002, India; Institute of computational Biology and Bioinformatics, Bioprudence Research Innovations LLP, Bhubaneswar, Odisha, Pin-751002, India; Department of Pharmacology, School of Pharmaceutical Sciences, Siksha O Anusandhan Deemed to be University, Bhubaneswar, Odisha, Pin-751003, India; Redox Biology and Proteomics Group, Department of Zoology, Ravenshaw Univarsity, Cuttack, Odisha, Pin-753003, India

**Keywords:** Diabetes mellitus, PPAR-γ, gamma-oryzanol, molecular docking, molecular dynamics simulation, scaffold analysis

## Abstract

Diabetes mellitus is one of the complex metabolic disorders associated with individuals leading sedentary lifestyles. It leads to several complications rendering the normal function of vital organs like heart, liver, kidney, eye and brain. Scientists and doctors across the globe are involved in research for understanding the complex genetics of this disorder and formulating newer therapeutics accordingly. The finding of potential chemical entities and their underlying agonists or antagonist activities significantly controls the disorder but with some consequences. Thus there is demand for natural compounds and indigenous treatment methods for controlling the disorder with least or no adverse consequences. In the current work we present computational prediction of gamma-oryzanol as potential agonist of human peroxisome proliferator-activated receptor gamma (PPAR-γ). A group of four gamma-oryzanol compound structures reported in PubChem database were downloaded and docked in the ligand binding site of five different human PPAR-γ structures reported in PDB database. It was observed that most of the gamma-oryzanol compounds occupied themselves in the ligand binding P1, P2, P3, P4 sites with similar orientations as that of co-crystal agonists. Their binding conformations were assisted by some reasonable docking scores (−7 to -11 kcal/mol) and hydrogen bond interactions with some important conserved amino acid residues lining the ligand binding site. Additionally we have done a comparative molecular dynamics studies to reveal the flexibility of gamma-oryzanol in the ligand binding site in comparison to the co-crystal agonist and a scaffold analysis using the structure of six agonists and gamma-oryzanol for fetching potential scaffolds which may helpful in designing of new chemical entities.

## 1. Introduction

Diabetes mellitus seems to remain as one of the major global public health concerns despite the implementation of several control measures either in the form of newer therapeutics or biomedical research. In recent decades the increase in prevalence of this metabolic disorder in several developing and developed countries is rendering their public health and socio-economic development [1-3]. The estimation of current status of diabetes mellitus by International Diabetes Federation (IDF) suggests that about 451 million cases were reported world wide in 2017 with a projection of 693 million cases by 2045 [4]. The brutality of this metabolic disorder is experienced as it accounts for over 80% of all premature non-communicable disease deaths along with cardiovascular, respiratory and cancer etc [5].

Current therapeutic management of this disorder includes lifestyle modification and pharmaceutical interventions [6, 7]. The lifestyle modification targets balance in energy by lowering caloric intake and enhancing expenditure by physical activities. The pharmaceutical interventions include administration of different drug molecules which target specific aspects of the disorder. The chronology of pharmaceutical intervention shows the use of insulin and insulin analogues, sulfonylureas, biguanides, alpha-glucosidase inhibitors, meglitinides, dipeptidylpeptidase-4 inhibitors and thiazolidinediones etc [8-10]. These anti diabetic agents interact with specific receptors or molecular targets expressed in diabetic conditions with the regulation of insulin sensitivity, insulin secretion, storage of fatty acids and glucose metabolism etc.

The human peroxisome proliferator-activated receptor (PPAR) is one of the most studied molecular targets for treatment of diabetes mellitus. It belongs to the steroid type nuclear receptor family and acts as ligand inducible transcription factor involved in various cellular processes [11-13]. Out of three reported isotypes of PPAR the PPAR-γ plays an important role in regulation of adipocyte differentiation, lipid metabolism, glucose homeostasis, inflammation and cell proliferation [14-16]. The activation of PPAR-γ is achieved by either endogenous ligands like fatty acids and prostanoids [17, 18] or synthetic thiazolidinedione agonists [19, 20], the former being weak agonists in comparison to the later as strong agonists. The thiazolidinedione series of compounds approved for treatment of diabetes mellitus were Troglitazone, Rosiglitazone and Pioglitazone with their specific roles of alleviating insulin resistance. But due to associated adverse side effects of these synthetic chemical agonists they were withdrawn from the market with restricted clinical application [21]. Thus there is a need for potential compounds that can be suitable agonists or partial agonists with improved glucose homeostasis activity and reduced side effects [22, 23].

Natural compounds isolated from different medicinal plants have been successfully used as drugs for the treatment of various diseases with least or no side effects [24]. These natural compounds are characterized by diverse chemical scaffolds optimized for performing different biological functions and act as important sources of drug discovery [25-28]. Gamma-oryzanol is the main bioactive compound isolated from bran layers of rice and used as potential anti-oxidative, anti-inflammatory and anti-cancer agents [29-31]. Recent investigations suggest that consumption of brown rice instead of white rice can improve the hyperglycemia in diabetic complications [32, 33]. Although gamma-oryzanol has a positive impact in the treatment of obesity related comorbidities [34-36] little is known about its action as potential agonist of PPAR-γ.

In the current investigation we target computational prediction of gamma-oryzanol as potential agonist of human PPAR-γ using the information availed by co-crystal agonists of PPAR-γ receptors reported in various public domains. The working hypothesis includes removal of reported co-crystal agonists from the ligand binding sites of PPAR-γ crystal structures and docking of gamma-oryzanol in the same site so as to reveal how efficiently it is occupied in the site in comparison to the co-crystal agonists assisted by the concepts of molecular docking, molecular dynamics simulations and scaffold analysis.

## 2. Materials and Methods

### 2.1 Materials

#### 2.1.1 Selection of human PPAR-γ receptor structures with co-crystal agonists

About six different crystal structures of human PPAR-γ receptor were searched in protein data bank (PDB, https://www.rcsb.org) [37] with their PDB IDs 1FM6, 1FM9, 1I7I, 1K74, 1NYX and 4PRG. These structures were reported to have potential co-crystal agonists with varied chemical compositions in the literature [38]. Thus the crystal structures of six receptors and the co-crytal agonists were downloaded in .pdb and .sdf format respectively for analysis. The pysiochemical analysis of these six agonists revealed that compounds Farglitazer, GW409544 and GW0072 have variationns from ideal properties (rules of five) than other three agonists Rosiglitazone, Tesaglitazar and Ragaglitazar as mentioned in Table-1.

**Table-1:** Important pysiochemical information of selected agonists

| Sl. No. | Agonist name | Agonist PDB ID | Receptor PDB ID | Receptor Chain | MW | XLogP | HBD | HBA | NR |
| --- | --- | --- | --- | --- | --- | --- | --- | --- | --- |
| 1 | Rosiglitazone | BRL | 1FM6 | D | 357.4 | 3.1 | 1 | 6 | 7 |
| 2 | Farglitazar | 570 | 1FM9 | D | 546.6 | 7.8 | 2 | 7 | 12 |
| 3 | Tesaglitazar | AZ2 | 1I7I | A | 408.5 | 3.5 | 1 | 7 | 11 |
| 4 | GW409544 | 544 | 1K74 | D | 510.6 | 6.8 | 2 | 7 | 12 |
| 5 | Ragaglitazar | DRF | 1NYX | A | 419.5 | 5 | 1 | 6 | 9 |
| 6 | GW0072 | 072 | 4PRG | A | 614.8 | 8.4 | 1 | 5 | 18 |

**MW:** Molecular Weight in Daltons, **XlogP:** Partition coefficient, **HBD:** Hydrogen Bond Donor count, **HBA:** Hydrogen Bond Acceptor count, **NR:** Number of Rotatable bonds

#### 2.1.2 Selection of Gamma-oryzanol type compounds from PubChem

The NCBI PubChem database (https://pubchem.ncbi.nlm.nih.gov/) [39] was searched with the keyword “gamma Oryzanol” resulting 8 compound entries, 65 substances entries, 141 literature entries and 77 patent entries. Out of eight compound entries the four compounds with PubChem CIDs (5282164, 51346127, 91971120, 134689750) were downloaded in 3D .sdf format and converted to .pdb format for further analysis. They were also renamed as compound1, compound2, compound3 and compound4 for ease in docking studies. All four compounds have similar physio-chemical properties (MW: 602.9, XlogP: 12.1, HBD: 1, HBA: 4 and NR: 9) with little variations in their structure conformations.

### 2.2 Methods

#### 2.2.1 Structural analysis and dissection of ligand binding site

All six selected PPAR-γ crystal structures were opened in PyMol [40] for their structural analysis. First they were structurally aligned with the calculations of root mean squared deviation (RMSD) values so as to reveal their structural relatedness. The highlighted RMSD values of the matrix shown in Table-2 predict structural resemblance between the crystal structures 1FM6-1FM9 (0.369), 1I7I-1K74 (0.517) and 1NYX-4PRG (0.737). Again for each analysis the ligand binding site was defined by selecting the residues lying 5Å of the included co-crystal agonists. In all six receptor structures the ligand binding site was defined by four designated subsites like P1, P2, P3 and P4 as shown in figure-1. The P1 site is defined by the carboxyl head group of the included co-crystal agonist. The P2 site is located slightly towards the right of the P1 site and both P3 and P4 sites are defined by the regions at the upper and lower distal ends. The amino acid residues included in the four subsites were P1 site (F282, C285, Q286, S289, H323, Y327, F363, H449, L469, Y473), P2 site (C285, R288, S289, I326, Y327, L330, F363, M364), P3 site (R288, E291, A292, E295, M329, E343) and P4 site (S255, E259, F264, H266, V277, A278, R280, I281, G284, F287, I341, S342, M348). A comparison of amino acid residues included among the four sites revealed that four residues (C285, S289, Y327 and F363) were common between P1 and P2 sites, one residue (R288) was common between P2 and P3 sites.

**Table-2:** Structural alignment of six PPAR-γ crystal structures showing the RMSD values.

|  | 1FM6 (A) | 1FM9 (D) | 1I7I (A) | 1K74 (D) | 1NYX (A) | 4PRG (A) |
| --- | --- | --- | --- | --- | --- | --- |
| 1FM6 (A) | 0.000 | **0.369** | 0.446 | 0.432 | 0.600 | 0.848 |
| 1FM9 (D) | **0.369** | 0.000 | 0.404 | 0.331 | 0.500 | 0.806 |
| 1I7I (A) | 0.446 | 0.404 | 0.000 | **0.517** | 0.396 | 0.701 |
| 1K74 (D) | 0.432 | 0.331 | **0.517** | 0.000 | 0.591 | 0.847 |
| 1NYX (A) | 0.600 | 0.500 | 0.396 | 0.591 | 0.000 | **0.737** |
| 4PRG (A) | 0.848 | 0.806 | 0.701 | 0.847 | **0.737** | 0.000 |

The RMSD values are presented in Å unit. The PPAR-γ crystal structure PDB ID and their corresponding chain information are presented in the first row and first column of the matrix. The diagonal values are 0 as per the definition of RMSD. The upper and lower triangles of the matrix represent the RMSD values of the two different structures.

**Figure-1:**
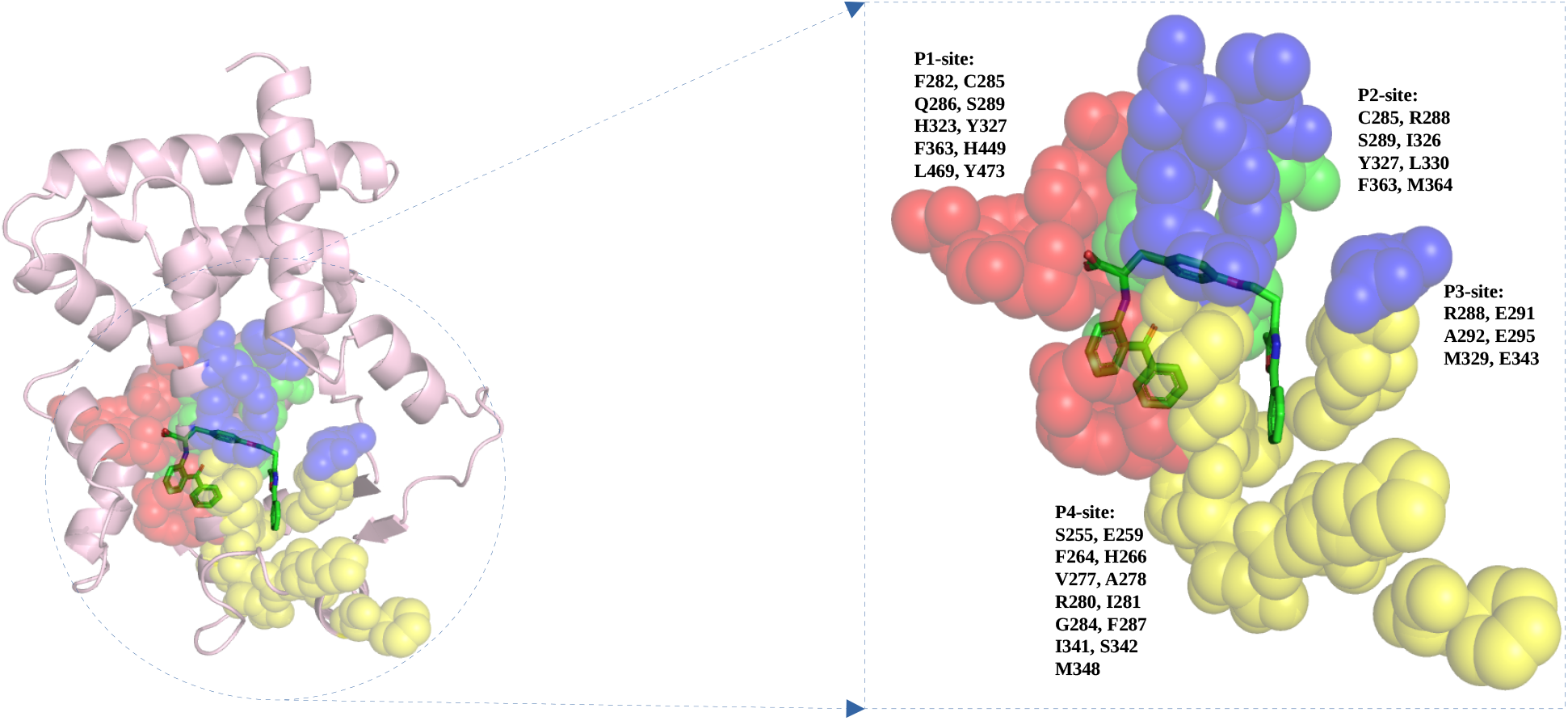
The ligand binding site of PPAR-receptor (PDB ID: 1FM6) with four subsites P1 site (red), P2 site (green), P3 site (blue) and P4 site (yellow) shown in sphere form. The amino acid residues comprising the four sites are shown adjacent to the sites.

#### 2.2.2 Preparation of selected receptor and grid boxes

All six PPAR-γ receptor structures were opened in AutoDockTools [41, 42] for preparation of grid boxes required for docking exercise. First the receptors were prepared by addition of polar hydrogen atoms and then addition of Kollman charges. Then the prepared receptors were saved in PDBQT format. The gridboxes were defined by selecting the co-crystal agonists and the grid parameters were saved after some modifications in X, Y and Z co-ordinate values.

#### 2.2.3 Preparation of gamma oryzanol compounds

All four gamma oryzanol compound structures were opened in AutoDockTools for their preparation. During preparation the compounds were added with Gasteiger charges and polar hydrogen atoms with merging of 57 non-polar hydrogens. About nine aromatic carbons were found per compound with the detection of ten rotatble bonds. Then the prepared compounds were saved in PDBQT format.

#### 2.2.4 Molecular docking of gamma oryzanol compounds on preprared receptors

All four prepared gamma oryzanol compounds were docked on five prepared receptor stuructures using AutoDock Vina [43, 44]. Prior to docking indivisual configuration files were prepared featuring the receptor name, ligand name, grid parameters, energy ranges and exhaustiveness etc. During docking exercise the receptor structure was fixed as rigid and the ligand molecules were made flexible so as to achieve suitable accommodation within the receptor structures.

#### 2.2.5 Molecular dynamics simulation studies

The dynamics of PPAR-γ receptor (4PRG chain A) and its complexes (complex1: 4PRG chain A with co-crystal agonist GW0072, complex2: 4PRG chain A with best docked pose of gamma-oryzanol compound2 exhibiting highest docking score) were studied by molecular dynamics (MD) simulations. The MD simulations were carried out in aqueous environment with Gromacs-2018.1 packages using amber99sb-ILDN force field [45]. The topology of the ligands (co-crystal agonist GW0072 and gamma-oryzanol compound2) was generated with the AM1-BCC charge model using Antechamber packages in AmberTools21 [46]. The PPAR-γ receptor alone and its complexes were solvated in triclinic boxes (to minimize the simulation box size) using TIP3P water model. The charges of each system (protein and its complexes) were neutralized by adding counter ions (sodium or chlorine) followed by energy minimization to remove the weak Van der Waals forces. The first equilibration (NVT equilibration) was performed at constant volume and temperature (300 K) using V-rescale thermostat for 1 ns [47]. The systems then minimized at constant pressure (1.0 bar) and temperature (300 K) for another 1 ns using Parrinello-Rahman barostat [48].The minimized structures were finally used to run 50 ns MD simulation in 10000 frames of each trajectory was saved. The trajectories were subjected to PBC corrections before analysis. All analysis was performed using standard gromacs utilities.

#### 2.2.6 Fetching of potential active scaffolds

Fetching of potential and active scaffolds among the chemical compounds seems to be an important step for the medicinal chemists involved in the design of new chemical entities. In order to fetch such active scaffolds among the six PPAR-γ agonists and gamma-oryzanol for design of potential agonist all six co-crystal agonists best docking poses and four best docking conformations of gamma-oryzanol were imported in scaffold hunter [49] in sdf format. During import the scaffold hunter automatically removed one compound (compound3_4prg_pose_1) based on the structure with the same SMILES being already imported. Then the nine compounds were assigned new properties like additional smiles, atom count calculator, DaylightBitFingerprinter, EStateBitFingerprinter, EStateNumericalFingerprinter and ExtendedConnectivityFingerprinter with the help of calculate properties facility. Once the dataset was assigned all calculated properties the scaffold tree was generated using the default ruleset.

## 3. Result and Discussions

### 3.1 Post docking analysis of gamma-oryzanol on six PPAR-γ receptor structure

About twenty four (four gamma-oryzanol compounds on six different PPAR-γ receptors) docking studies were analysed by opening the receptor structures first and then loading the docking conformations. The docking conformations with the best docking scores were first selected from all twenty four docking studies and tabulated (Table-3) for comparative analysis. From the table it was quite evident that the docking scores were reasonable (−6.9 kcal/mol to -11.8 kcal/mol) on which further analysis can be done.

**Table-3:** Post docking analysis of best docking scores for all twenty four docking

| Sl. No. | 1FM6 | 1FM9 | 1I7I | 1K74 | 1NYX | 4PRG |
| --- | --- | --- | --- | --- | --- | --- |
| Compound1 | -9.6 | -8.9 | -9.7 | -7.0 | -9.0 | -11.4 |
| Compound2 | -10.1 | -9.2 | -9.5 | -7.0 | -8.6 | -11.8 |
| Compound3 | -8.8 | -8.7 | -9.4 | -7.5 | -9.0 | -10.7 |
| Compound4 | -9.1 | -9.5 | -9.0 | -6.9 | -8.5 | -9.9 |

The docking scores associated with the best docking conformation is not alone sufficient for gathering meaningful information; rather it has to reflect the number of ideal hydrogen bond interactions between the residue atom and ligand atom. Thus all nine docking conformations for twenty four docking studies were further analysed by selecting the residues within the 5 Å area of gamma-oryzanol compounds.

The post docking analysis of gamma-oryzanol compounds on the first set of PPAR-γ receptors (PDB ID: 1FM6 and 1FM9) is summarized in Table-4. The residues of 1FM6 involved in hydrogen bond interaction with the gamma-oryzanol compounds are L288, R230, R288, Q294, Y327, S342, Q345, H466 and H499 from which Y327 and H449 lie in P1 site, R288 and Y327 lie in P2 site, R288 lies in P3 site and S342 lies in P4 site. Similarly the residues of 1FM9 involved in hydrogen bond interaction with the gamma-oryzanol compounds are L228, L255, E259, Q273, Q283, Q286, R288, L340, S342, Q345, H449 and S464 from which Q286 and H449 lie in P1 site, R288 shares equally in P2 and P3 sites and E259 and S342 lie in P4 site. This short of interaction suggests good orientation of gamma-oryzanol compounds in the ligand binding site as that of co-crystal agonist Rosiglitazone of 1FM6 and Farglitazar of 1FM9 with some reasonable docking score as mentioned in Table-4.

**Table-4:**
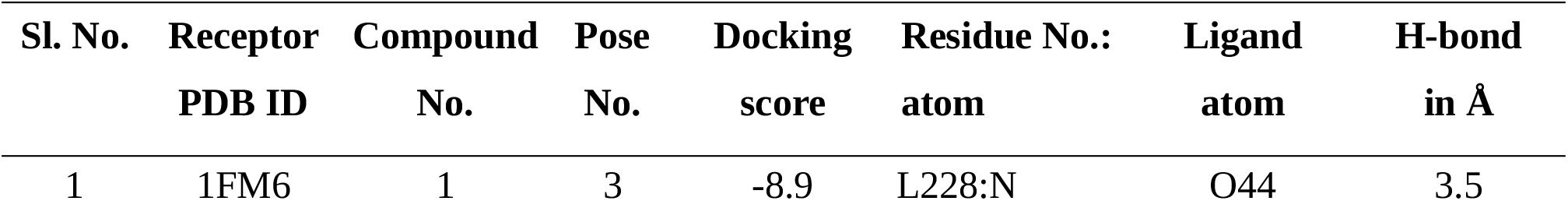
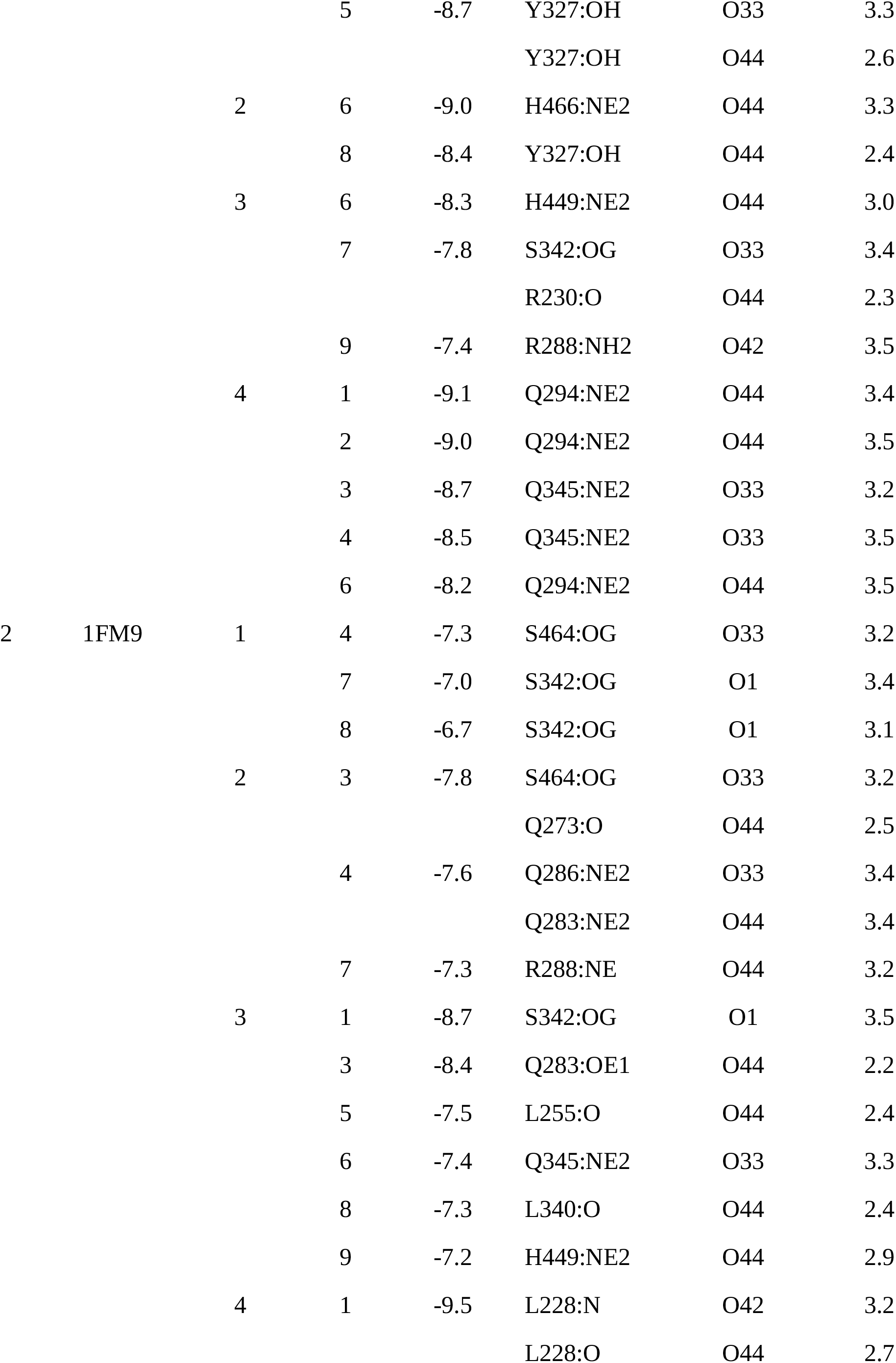
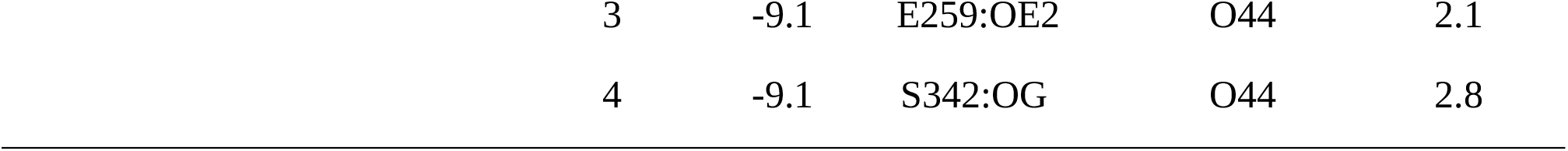
Post docking interaction analysis for receptors 1FM6 and 1FM9

The post docking analysis of gamma-oryzanol compounds on the second set of PPAR-γ receptors (PDB ID: 1I7I and 1K74) is summarized in Table-5. The residues of 1I7I involved in hydrogen bond interaction with the gamma-oryzanol compounds are K263, S289, S342, K343, H449, S464 and H466 from which S289 and H449 lie in P1 site, S289 lies in P2 site and S342 lies in P4 site. None of the interacting residues lie in P3 site. Similarly the residues of 1K74 involved in hydrogen bond interaction with the gamma-oryzanol compounds are L228, E272, S274, Q283, Q286, E291, S464 and H466 from which Q286 lies in P1 site and E291 lies in P3 site. None of the interacting residues lie in P2 site and P3 site. This short of interaction suggests less good orientation of gamma-oryzanol compounds in the ligand binding site as that of co-crystal agonist Tesaglitazar of 1I7I and GW409544 of 1K74 with some reasonable docking score as mentioned in Table-5.

**Table-5:** Post docking interaction analysis for receptors 1I7I and 1K74

| Sl. No. | Receptor<br>PDB ID | Compound<br>No. | Pose<br>No. | Docking<br>score | Residue No.:<br>atom | Ligand<br>atom | H-bond<br>in Å |
| --- | --- | --- | --- | --- | --- | --- | --- |
| 1 | 1I7I | 1 | 4 | -8.3 | S342:OG | O33 | 3.4 |
|  |  |  | 9 | -8.0 | H449:NE2 | O44 | 3.1 |
|  |  | 2 | 4 | -8.6 | S342:OG | O33 | 3.4 |
|  |  |  | 5 | -8.5 | S289:OG | O44 | 2.4 |
|  |  |  | 7 | -8.2 | S289:OG | O44 | 2.7 |
|  |  |  | 8 | -8.1 | H466:N | O44 | 3.0 |
|  |  | 3 | 3 | -8.4 | K263:NZ | O1 | 3.5 |
|  |  |  |  |  | K343:N | O44 | 3.4 |
|  |  |  | 9 | -7.9 | S464:O | O44 | 1.9 |
|  |  |  |  |  | H466:N | O44 | 3.1 |
|  |  | 4 | 3 | -8.7 | S342:N | O33 | 3.1 |
|  |  |  | 7 | -8.2 | S464:O | O44 | 2.6 |
|  |  |  |  |  | H466:N | O44 | 3.0 |
| 2 | 1K74 | 1 | 2 | -6.9 | E291:OE2 | O44 | 2.0 |
|  |  |  | 4 | -6.5 | Q286:NE2 | O44 | 3.4 |
|  |  |  |  |  | S464:O | O44 | 2.3 |
|  |  |  | 7 | -6.3 | Q283:NE2 | O1 | 3.5 |
|  |  |  |  |  | S274:OG | O44 | 2.8 |
|  |  | 2 | 2 | -6.7 | Q283:NE2 | O1 | 3.4 |
|  |  |  | 4 | -6.6 | E291:OE2 | O44 | 2.0 |
|  |  |  | 9 | -6.2 | Q286:NE2 | O44 | 3.4 |
|  |  | 3 | 1 | -7.5 | Q286:NE2 | O44 | 3.3 |
|  |  |  | 4 | -6.8 | Q286:OE1 | O44 | 2.5 |
|  |  |  |  |  | Q286:NE2 | O44 | 3.5 |
|  |  |  | 5 | -6.7 | Q286:NE2 | O42 | 3.3 |
|  |  |  |  |  | Q286:NE2 | O44 | 3.3 |
|  |  |  |  |  | S464:O | O44 | 2.6 |
|  |  |  | 6 | -6.4 | E272:N | O44 | 3.3 |
|  |  | 4 | 2 | -6.9 | Q283:NE2 | O42 | 3.4 |
|  |  |  |  |  | Q283:NE2 | O44 | 3.5 |
|  |  |  |  |  | Q286:NE2 | O44 | 3.4 |
|  |  |  | 4 | -6.7 | H466:ND1 | O44 | 2.7 |
|  |  |  | 9 | -5.8 | L228:O | O44 | 2.3 |

The post docking analysis of gamma-oryzanol compounds on the third set of PPAR-γ receptors (PDB ID: 1NYX and 4PRG) is summarized in Table-6. The residues of 1NYX involved in hydogen bond interaction with the gamma-oryzanol compounds are R288, L340, S342, S464 and H466 from which none of the interacting residue lies in P1 site, R288 is shared by both P2 site and P3 site and S342 lies in P4 site. Similarly the residues of 4PRG involved in hydrogen bond interaction with the gamma-oryzanol compounds are L228, K263, K265 and E295 from which only E295 lies in the P3 site and none of the interacting residues lie in P1, P2 and P4 sites. This short of interaction suggests good orientation of gamma-oryzanol compounds in the ligand binding site as that of co-crystal agonist Ragaglitazar of 1NYX and GW0072 of 4PRG with some reasonable docking score as mentioned in Table-6.

**Table-6:**
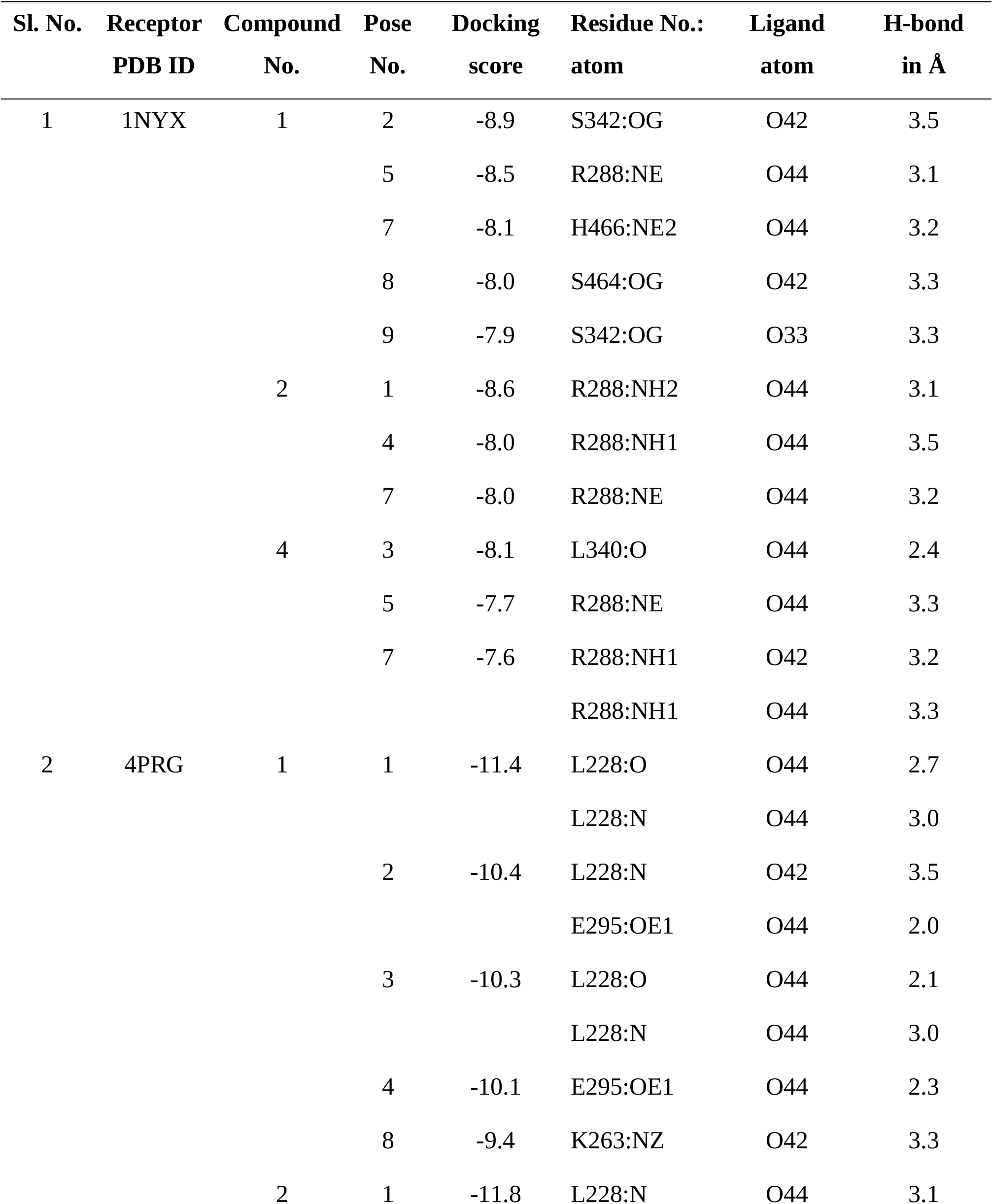
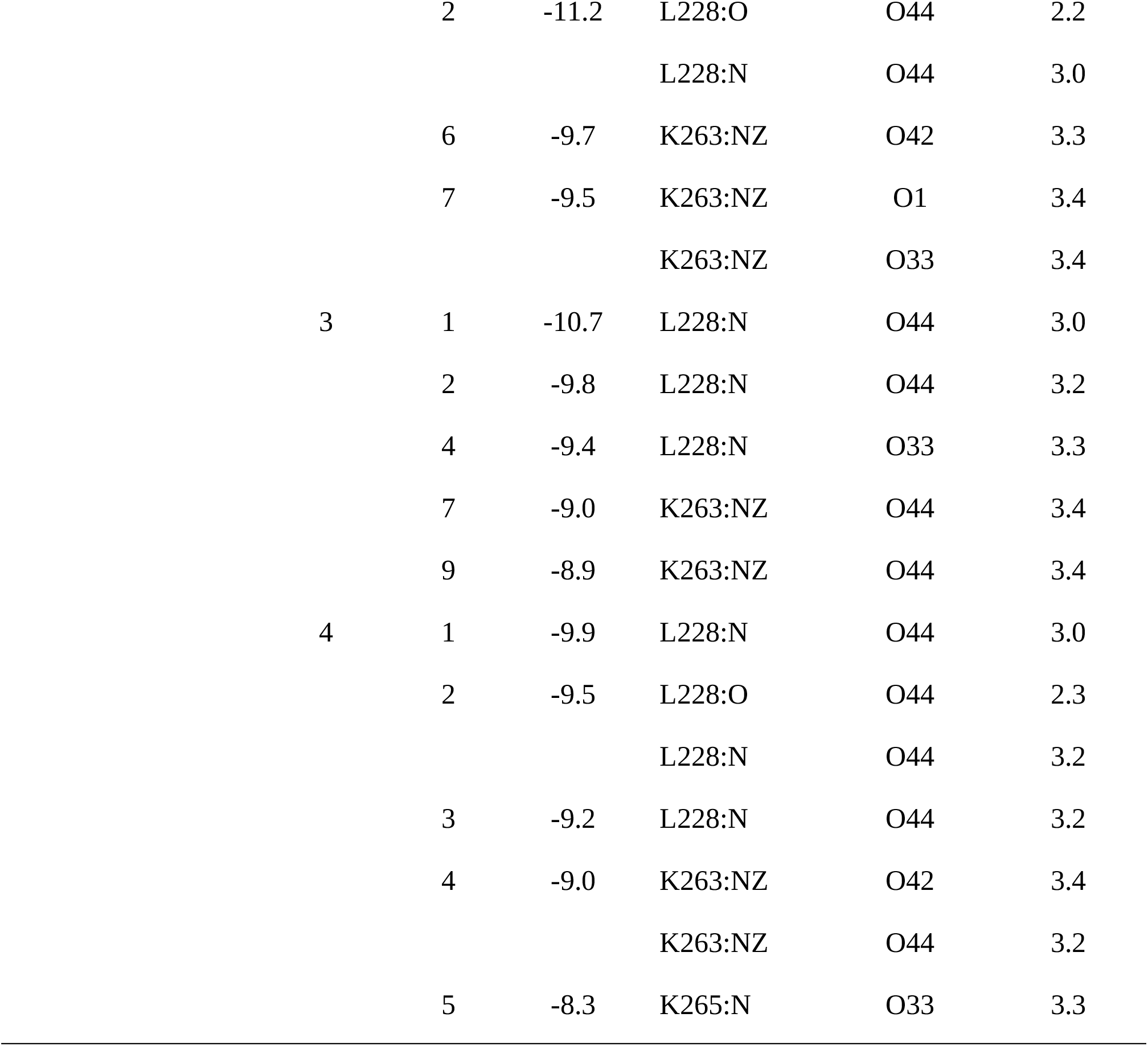
Post docking interaction analysis for receptors 1NYX and 4PRG

| Sl. No. | Receptor<br>PDB ID | Compound<br>No. | Pose<br>No. | Docking<br>score | Residue No.:<br>atom | Ligand<br>atom | H-bond<br>in Å |
| --- | --- | --- | --- | --- | --- | --- | --- |
| 1 | 1NYX | 1 | 2 | -8.9 | S342:OG | O42 | 3.5 |
|  |  |  | 5 | -8.5 | R288:NE | O44 | 3.1 |
|  |  |  | 7 | -8.1 | H466:NE2 | O44 | 3.2 |
|  |  |  | 8 | -8.0 | S464:OG | O42 | 3.3 |
|  |  |  | 9 | -7.9 | S342:OG | O33 | 3.3 |
|  |  | 2 | 1 | -8.6 | R288:NH2 | O44 | 3.1 |
|  |  |  | 4 | -8.0 | R288:NH1 | O44 | 3.5 |
|  |  |  | 7 | -8.0 | R288:NE | O44 | 3.2 |
|  |  | 4 | 3 | -8.1 | L340:O | O44 | 2.4 |
|  |  |  | 5 | -7.7 | R288:NE | O44 | 3.3 |
|  |  |  | 7 | -7.6 | R288:NH1 | O42 | 3.2 |
|  |  |  |  |  | R288:NH1 | O44 | 3.3 |
| 2 | 4PRG | 1 | 1 | -11.4 | L228:O | O44 | 2.7 |
|  |  |  |  |  | L228:N | O44 | 3.0 |
|  |  |  | 2 | -10.4 | L228:N | O42 | 3.5 |
|  |  |  |  |  | E295:OE1 | O44 | 2.0 |
|  |  |  | 3 | -10.3 | L228:O | O44 | 2.1 |
|  |  |  |  |  | L228:N | O44 | 3.0 |
|  |  |  | 4 | -10.1 | E295:OE1 | O44 | 2.3 |
|  |  |  | 8 | -9.4 | K263:NZ | O42 | 3.3 |
|  |  | 2 | 1 | -11.8 | L228:N | O44 | 3.1 |

|  |  |  |  |  |  |
| --- | --- | --- | --- | --- | --- |
|  | 2 | -11.2 | L228:O | O44 | 2.2 |
|  |  |  | L228:N | O44 | 3.0 |
|  | 6 | -9.7 | K263:NZ | O42 | 3.3 |
|  | 7 | -9.5 | K263:NZ | O1 | 3.4 |
|  |  |  | K263:NZ | O33 | 3.4 |
| 3 | 1 | -10.7 | L228:N | O44 | 3.0 |
|  | 2 | -9.8 | L228:N | O44 | 3.2 |
|  | 4 | -9.4 | L228:N | O33 | 3.3 |
|  | 7 | -9.0 | K263:NZ | O44 | 3.4 |
|  | 9 | -8.9 | K263:NZ | O44 | 3.4 |
| 4 | 1 | -9.9 | L228:N | O44 | 3.0 |
|  | 2 | -9.5 | L228:O | O44 | 2.3 |
|  |  |  | L228:N | O44 | 3.2 |
|  | 3 | -9.2 | L228:N | O44 | 3.2 |
|  | 4 | -9.0 | K263:NZ | O42 | 3.4 |
|  |  |  | K263:NZ | O44 | 3.2 |
|  | 5 | -8.3 | K265:N | O33 | 3.3 |

### 3.2 Analysis of molecular dynamics simulations

The primary analysis of molecular dynamics simulation of PPAR-γ receptor and two complexes under aqueous conditions was done by calculation of root mean squared deviation (RMSD) and root mean square fluctuation (RMSF) of Cα atoms each residue of each system. The analysis of RMSD plot of PPAR-γ and its complexes revealed that three systems quickly stabilized after 5 ns as RMSD remaining below 0.4 nm throughout the simulation (Figure-2A). The average RMSD of each system was found to be less than 0.32 nm, thus indicating a good stability of the systems. The analysis of RMSF plot of PPAR-γ and its complexes revealed that of most amino acid residues of PPAR-γ and its complexes had RMSF values below 0.4 nm (Figure-2B) which suggests their stability in aqueous environment. Further analysis of simulation was done by assessment of radius of gyration (Rg) and solvent accessible surface area (SASA). The Rg is an important parameter in measuring the compactness of a protein structure and SASA of macromolecules is considered as a decisive parameter in predicting their folding and stability. The analysis of Rg plot revealed that the Rg of all three systems remained constant throughout the simulation time (Figure-2C) thus predicting their stability. The analysis of SASA plot revealed that the average SASA of three systems were 215.43, 217.55 and 213.97 nm2 respectively (Figure-2D) which also suggests their stability in aqueous conditions.

**Figure-2:**
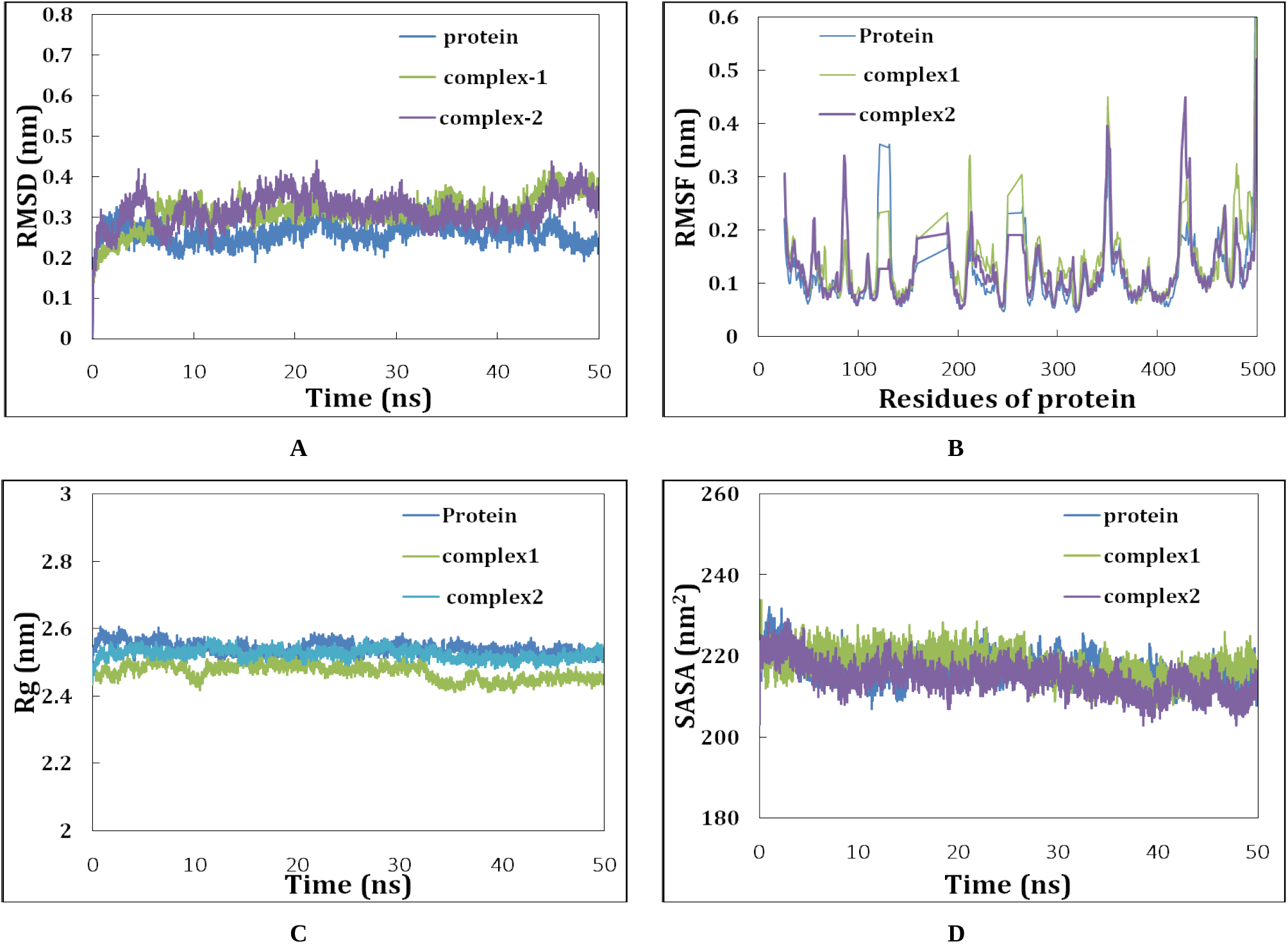
Analaysis of molecular dynamics simulations of PPAR-γ receptor and two complexes, A: RMSD plot, **B:** RMSF plot, **C:** Rg plot and **D:** SASA plot.

### 3.3 Analysis of potential scaffolds

After scaffold analysis a scaffold tree view was generated with molecules being sorted by largest fragment of DaylightBitFingerprinter and the corresponding scaffolds being sorted by number of aromatic rings (SCPnoAroRings). About four (O1CNCC1, C1CCCCC1, S1CNCC1 and O1CCNCC1) potential scaffolds were generated from decomposition of nine compounds as summarized in Table-7. The first scaffold (C1CCCCC1) was represented by about five compounds like GW0072 (Thiazolidinone derivative), Tesaglitazer (dihydrocinnamate derivative) and three gamma-oryzanol compounds. The second scaffold (O1CNCC1) was represented by only two compounds like GW0409544 (L-tyrosine derivative) and Farglitazer (L-tyrosine derivative). The third (S1CNCC1) and fourth (O1CCNCC1) scaffold were unique and represented by potential agonists like Rosiglitazone (thiazolidinediones derivative) and Ragaglitazer (n-substituted phenoxazines) respectively.

**Table-7:**
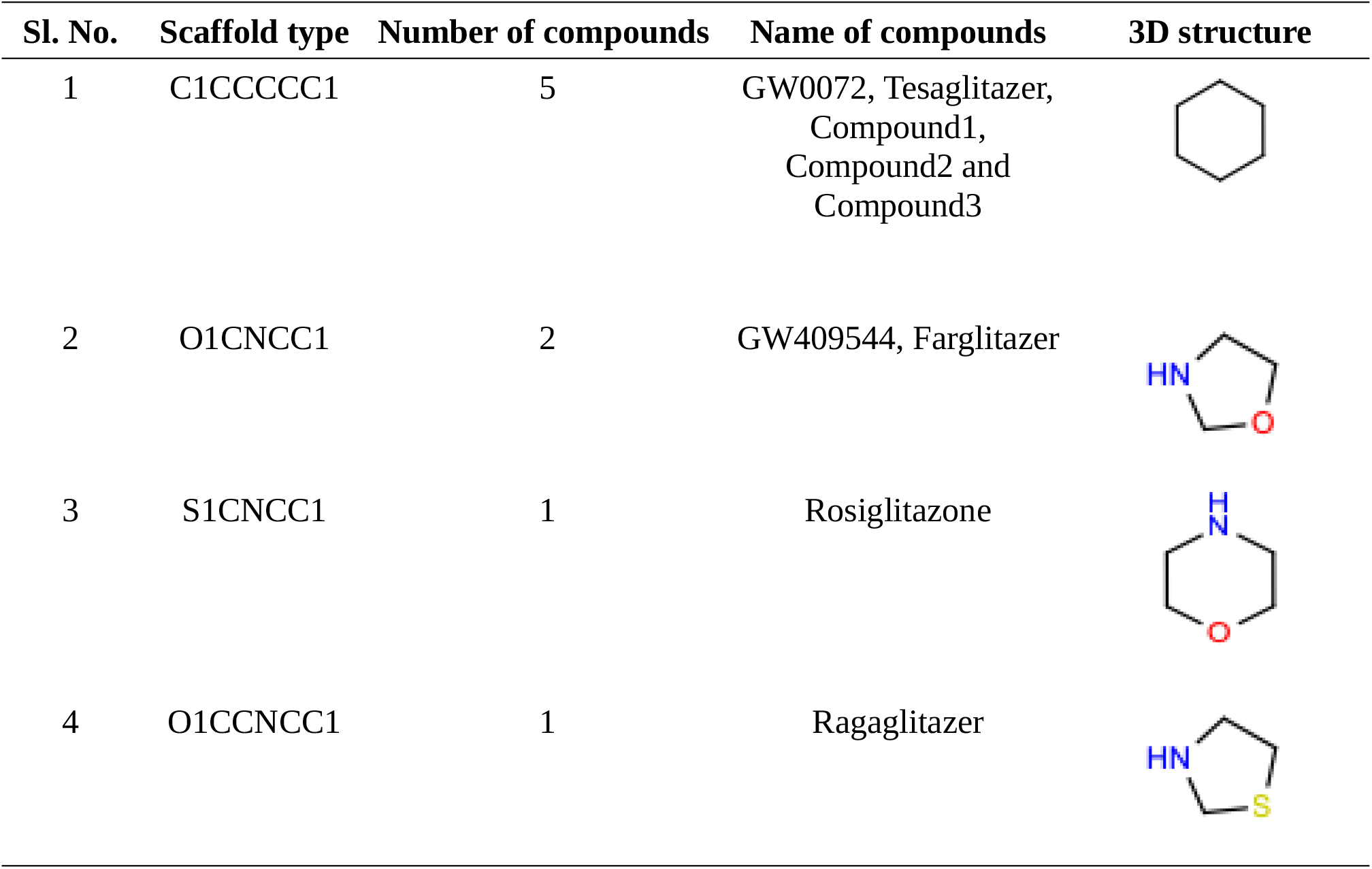
Summary of scaffold analysis

| Sl. No. | Scaffold type | Number of compounds | Name of compounds | 3D structure |
| --- | --- | --- | --- | --- |
| 1 | C1CCCCC1 | 5 | GW0072, Tesaglitazer, Compound1, Compound2 and Compound3 |  |
| 2 | O1CNCC1 | 2 | GW409544, Farglitazer |  |
| 3 | S1CNCC1 | 1 | Rosiglitazone |  |
| 4 | O1CCNCC1 | 1 | Ragaglitazer |  |

### 3.4 Gamma-oryzanol as potential agaonist of PPAR-γ

#### 3.4.1 Gamma-oryzanol accommodates in the ligand binding site of PPAR-γ

As per our working hypothesis we have removed the reported co-crystal agonists from six different PPAR-γ receptors and docked four gamma-oryzanol compounds in their ligand binding sites. In all six cases it was observed that gamma-oryzanol suitbaly accommodates in the ligand binding site with some variations in their orientations which may be due selection of ten rotatable bonds (torsdof=10) during ligand preparation time. The variations observed were also different for all three sets of PPAR-γ receptors under study.

The first set of receptors 1FM6 and 1FM9 have co-crystal agonists like Rosiglitazone and Farglitazar respectively. In 1FM6 it was observed that the main 5-methyl-1,3-thiazolidine-2,4-dione scaffold of Rosiglitazone goes deep into the P1 site, the middle 1-methoxy-4-methylbenzene scaffold remains in between P2 and P3 sites and the last N,N-dimethylpyridin-2-amine scaffold remains in the P4 site. It shows hydrogen bond interactions with residues S289, H323 and Y473 of P1 site. In comparison to Rosiglitazone the ferulate scaffold of gamma-oryzanol remains in between P2 and P3 sites and the remaining steroidal scaffold goes deep in the P4 site as shown in Figure-3A. It shows hydrogen bond interactions with residues Y327 and H449 of P1 site, R288 and Y327 of P2 site, R288 of P3 site and S342 of P4 site. In 1FM9 it was observed that the diphenylmethanone scaffold of Farglitazar goes deep into the P1 and P2 sites, the middle 1-methoxy-4-methylbenzene scaffold remains in the P2 site and the last 5-Methyl-2-phenyloxazole scaffold remains in the P4 site. It shows hydrogen bond interactions with residues S289, H323, Y473 and H449 of P1 site. In comparison to Farglitazar the ferulate scaffold of gamma-oryzanol goes deep into the P4 site and the remaining steroidal scaffold remains in between P1 and P2 sites as shown in Figure-3B. It shows hydrogen bond interactions with residues Q286 and H449 of P1 site, R288 of P2 site and E259 and S342 of P4 site.

**Figure-3:**
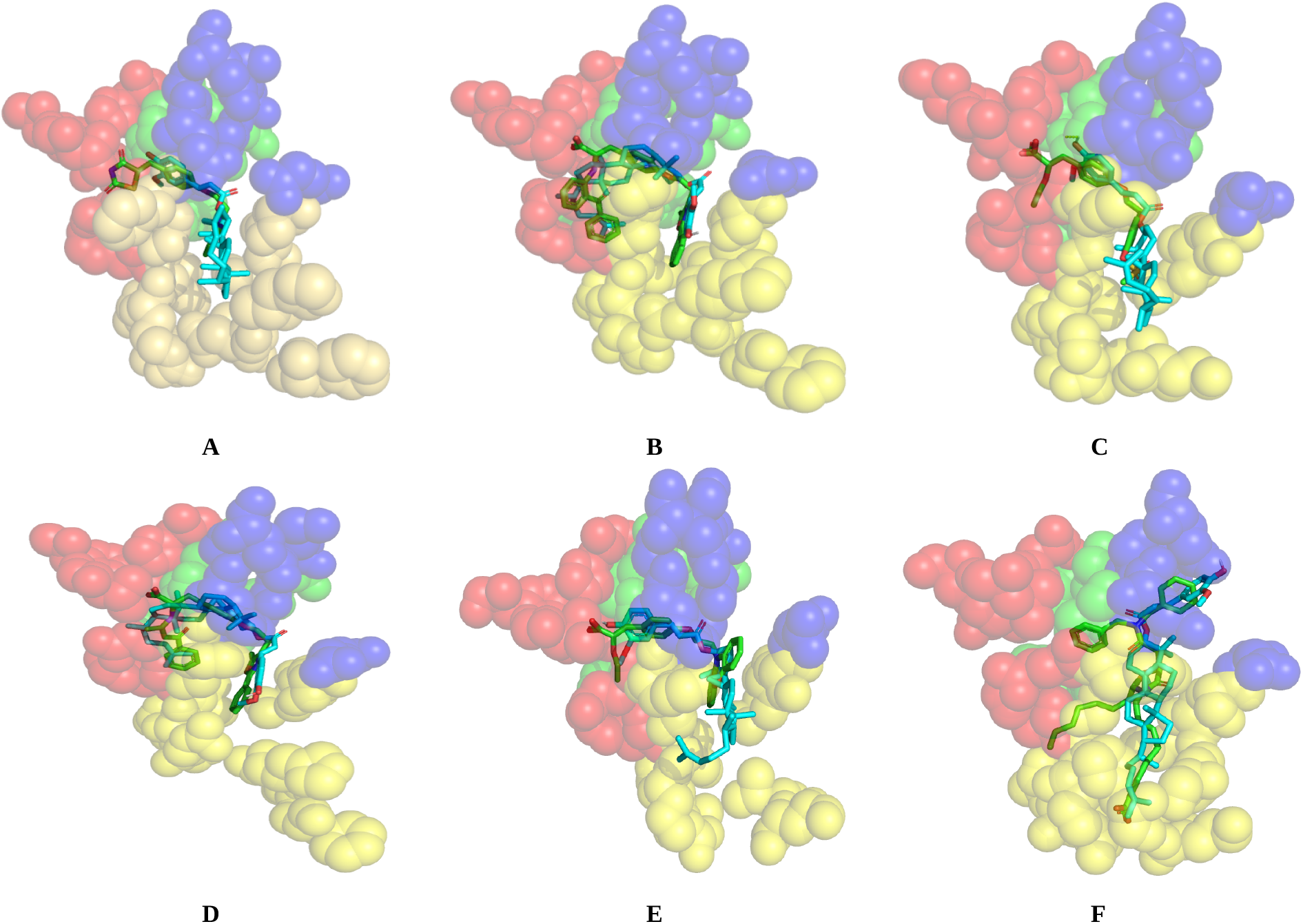
The superposition of gamma-oryzanol and the co-crystal agonist of the PPAR-γ receptors in the ligand binding site with four subsites P1 site (red), P2 site (green), P3 site (blue) and P4 site (yellow) shown in sphere form. **A:** gamma-oryzanol and Rosiglitazone **B:** gamma-oryzanol and Farglitazar **C:** gamma-oryzanol and Tesaglitazar **D:** gamma-oryzanol and GW409544 **E:** gamma-oryzanol and Ragaglitazar **F:** gamma-oryzanol and GW0072.

The second set of receptors 1I7I and 1K74 have co-crystal agonists like Tesaglitazar and GW409544 respectively. In 1I7I it was observed that the 2-Ethoxy-3-(4-methylphenyl)propanoic acid scaffold remains in between P1 and P2 sites and the methylsulfoxy scaffold goes deep into the P4 site. It shows hydrogen bond interactions with residues S289, H323, Y473 and H449 of P1 site. In comparison to Tesaglitazar the ferulate scaffold of gamma-oryzanol remains in between P2 and P1 sites and the remaining steroidal scaffold goes deep in the P4 site as shown in Figure-3C. It shows hydrogen bond interactions with residues S289 and H449 of P1 site and S342 of P4 site. In 1K74 it was observed that the 1-phenylethanone scaffold of GW409544 goes deep into the P1 site, the middle 1-methoxy-4-methylbenzene scaffold remains in between the P2 and P3 site and the last 5-Methyl-2-phenyloxazole scaffold remains in the P4 site. It shows hydrogen bond interactions with residues S289, H323 and H449 of P1 site. In comparison to GW409544 the steroidal scaffold remains in between P1 and P2 sites and the remaining ferulate scaffold goes deep into the P4 site as shown in Figure-3D. It shows hydrogen bond interactions with residues Q286 of P1 site and E291 of P3 site

The third set of receptors 1NYX and 4PRG have co-crystal agonists like Ragaglitazar and GW0072 respectively. In 1NYX it was observed that the 2-ethoxy-3-phenylpropanoic acid scaffold of Ragaglitazar remains in between P1 and P2 sites with the carboxyl group deep into the P1 site and the Phenoxazine scaffold remains in between the P3 and P4 sites. It shows hydrogen bond interactions with residues H323, H449 and Y473 of P1 site. In comparison to Ragaglitazar the ferulate scaffold of gamma-oryzanol remains in between P1 and P2 sites and the steroidal scaffold remains in between P3 and P4 sites as shown in Figure-3E. It shows hydrogen bond interactions with residues R288 of P2 site and S342 of P4 site. In 4PRG it was observed that the dibenzylamide scaffold of GW0072 remains within P2 and P3 sites, the carboxyl goes deep into the P4site and the thiazolidinone scaffold remains in between P3 and P4 sites. It shows hydrogen bond interactions with residues S288 of P3 site and R280 and S342 of P4 site. In comparison to GW0072 the ferulate scaffold of gamma-oryzanol remains in the P3 site and steroidal scaffold goes deep into the P4 site as shown in Figure-3F. It shows hydrogen bond interactions with residues E295 of P3 site.

#### 3.4.2 Gamma-oryzanol exhibits flexibility while occupied in ligand binding site

The post docking analysis of gamma-oryzanol compound2 on PPAR-γ receptor 4PRG was compared with the co-crystal agonist GW0072 which revealed greater flexibility of the GW0072 as compared to the gamma-oryzanol compound2 in the ligand binding site (Figure-4A). This sort of flexibility is achieved by the presence of about eighteen rotatable bonds in GW0072 as compared to only ten rotatable bonds of gamma-oryzanol compounds. Again for confirming the greater flexibility of GW0072 in comparision to gamma-oryzanol compound the RMSF of individual atoms of both GW0072 and gamma-oryzanol compound2 as complexed with the PPAR-γ receptor 4PRG proteins were also calculated from the simulation study. The RMSF plot shown in Figure-4B shows some variations of RMSF values of the atoms indicating the dynamical shift of the compounds under study. It also suggests that the RMSF of compound1 (GW0072) is more than the compound2 (gamma-oryzanol compound2) which makes it more flexible in the ligand binding site.

**Figure-4:**
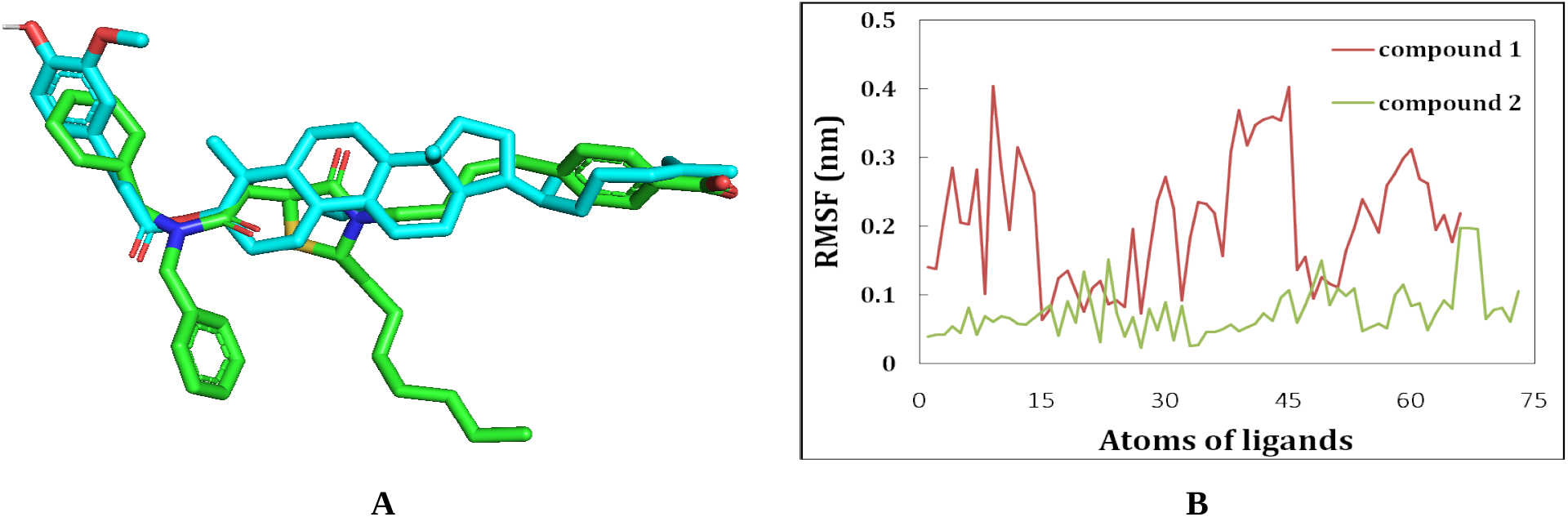
**A:** The superposition of best docked conformation of gamma-oryzanol (cyan colouration) and the co-crystal agonist GW0072 (green colouration) of PPAR-γ receptor (PDB ID: 4PRG) **B:** The RMSF of individual atoms of compound1 (GW0072) and compound2 (gamma-oryzanol) when complexed with the PPAR-γ receptor (PDB ID: 4PRG)

#### 3.4.3 Gamma-oryzanol possess same chemical scaffolds as that of designated agonists

The first level of hierarchy of the scaffold tree represents the four potential scaffolds decomposed from nine compounds. The second level of hierarchy represents the two member scaffolds as shown in Figure-5A and subsequent new scaffolds are added with the increase in the hierarchy. The hierarchy of scaffolds stops with the inclusion of the last scaffold thus completing the chemical structure of the compounds under study. In the scaffold decomposition process the compounds under study are decomposed in the reverse way upto the hierarchy with single member scaffolds. It was observed that Gamma-oryzanol goes through five stages of decomposition to reach its major scaffold C1CCCCC1. In comparison Tesaglitazer goes through only one stage of decomposition and GW0072 goes through three stages of decomposition to reach the same scaffold C1CCCCC1. Thus in our study targeting gamma-oryzanol as a suitable agonist of PPAR-γ receptor it was observed that it possesses the same potential scaffold as that of some designated agonists like GW0072 and Tesaglitazar (Figure-5B).

**Figure-5:**
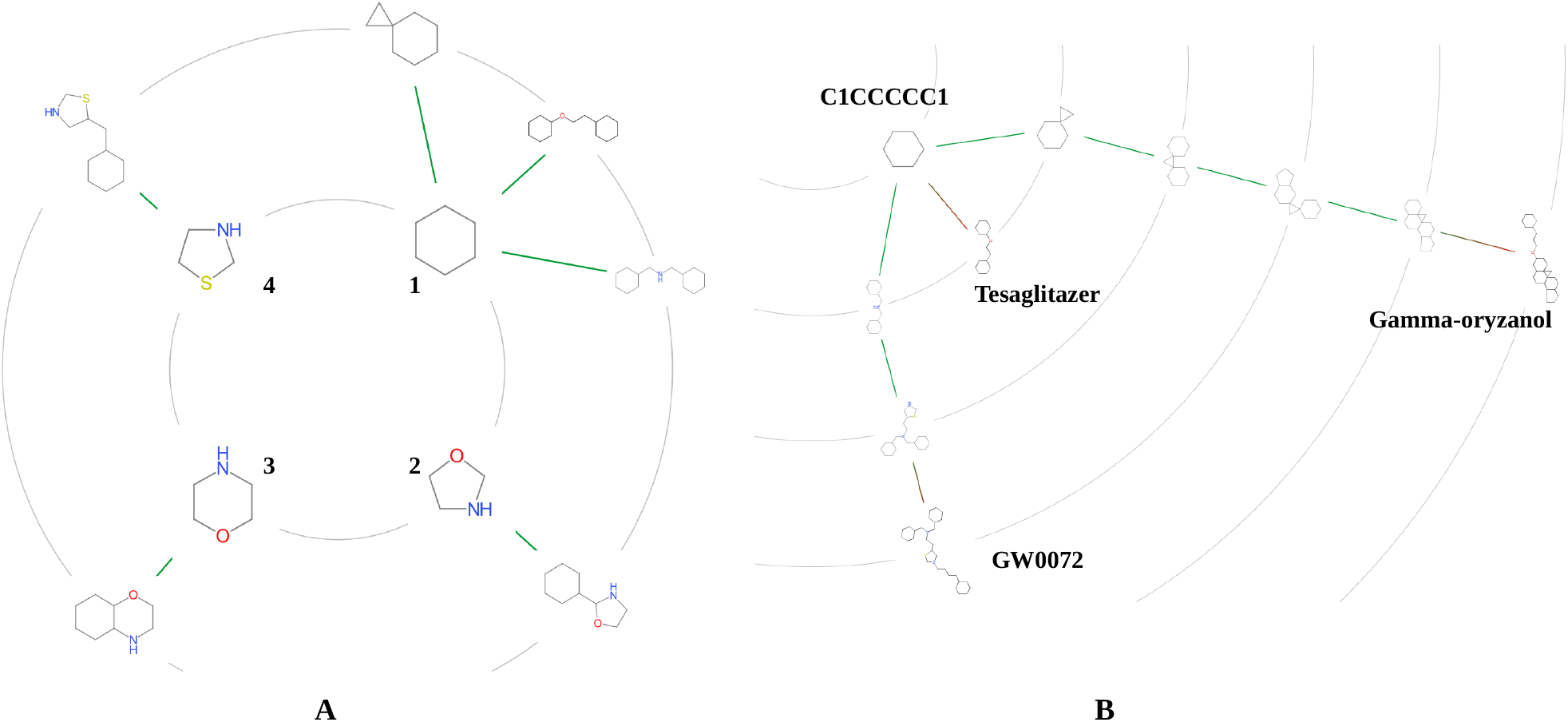
Graphical representation of potential scaffolds occupied in the gamma-oryzanol compounds and six co-crystal agonists. **A:** The scaffold tree showing the first hierarchy of four potential scaffolds and second hierarchy of two memebered scaffolds **B:** The scaffold decomposition process of first scaffold C1CCCCC1 from its chemical compounds.

## 4. Conclusions

In the process of computational prediction of gamma-oryzanol as potential agonist of human PPAR-γ receptor we have docked four gamma-oryzanol compounds in the ligand binding sites of six human PPAR-γ receptors complexed with reported potential agonists. The post docking analysis revealed that gamma-oryzanol suitably binds in the ligand binding sites of the six receptors under study with a similar shot of orientation as that of the complexed agonists. A comparative molecular dynamics study of the highly scored (−11.8 kcal/mol) gamma-oryzanol compound and PPAR-γ receptor complex with reported co-crystal agonist and the same PPAR-γ receptor complex also revealed suitable accommodation of gamma-oryzanol in the ligand binding site. At last fetching of potential active scaffolds among the highly scored gamma-oryzanol compounds and all six co-crystal agonists also revealed that gamma-oryzanol possesses the same active scaffold as possessed by some reported agonists. Our computational prediction may be useful in the design of new chemical entities with some modification of gamma-oryzanol moiety that can act as suitable agonists of the human PPAR-γ receptor.

## 5. Acknowledgements

The authors are grateful to The Principal of Dadhichi colllege of Pharmacy, Faculty members of SOA University, Ravenshaw University and Bioprudence Research Innovations LLP for providing essential facilities in conducting research leading to the level of publications. The award of ICMR-Research Assocateship (No.3/1/3(5)/Endo-fellowship/22-NCD-III) for Pabitra Mohan Behera is duly acknowledged.

## References

1. Patterson, C. C. et al. Trends and cyclical variation in the incidence of childhood type 1 diabetes in 26 European centres in the 25 year period 1989–2013: a multicentre prospective registration study. Diabetologia 62, 408–417 (2019).

2. Wang, L. et al. Prevalence and ethnic pattern of diabetes and prediabetes in China in 2013. JAMA 317, 2515–2523 (2017).

3. Dwyer-Lindgren, L., Mackenbach, J. P., Van Lenthe, F. J., Flaxman, A. D. & Mokdad, A. H. Diagnosed and undiagnosed diabetes prevalence by County in the U.S., 1999–2012. Diabetes Care 39, 1556–1562 (2016).

4. Cho, N. H. et al. IDF Diabetes atlas: global estimates of diabetes prevalence for 2017 and projections for 2045. Diabetes Res. Clin. Pract. 138, 271–281 (2018).

5. Forouzanfar, M. H. et al. Global, regional, and national comparative risk assessment of 79 behavioural, environmental and occupational, and metabolic risks or clusters of risks, 1990–2015: a systematic analysis for the Global Burden of Disease Study 2015. Lancet 388, 1659–1724 (2016).

6. Ford ES, Li C, Zhao G. Prevalence and correlates of metabolic syndrome based on a harmonious definition among adults in the US. J Diabetes 2010;2: 180–93.

7. Grundy SM, Brewer Jr HB, Cleeman JI, Smith Jr SC, Lenfant C. Definition of metabolic syndrome: report of the National Heart, Lung, and Blood Institute/American Heart Association conference on scientific issues related to definition. Circulation 2004;109:433–8.

8. Berenson DF, Weiss AR, Wan ZL, Weiss MA. Insulin analogs for the treatment of diabetes mellitus: therapeutic applications of protein engineering. Ann NY Acad Sci 2011;1243:E40–54.

9. Mizuno CS, Chittiboyina AG, Kurtz TW, Pershadsingh HA. Avery MA. Type 2 diabetes and oral antihyperglycemic drugs. Curr Med Chem 2008;15: 61–74.

10. Seino S, Takahashi H, Takahashi T, Shibasaki T. Treating diabetes today: a matter of selectivity of sulphonylureas. Diabetes Obes Metab 2012;14(Suppl1):9–13.

11. Fox CS, Pencina MJ, Meigs JB, et al. Trends in the incidence of type 2 diabetes mellitus from the 1970s to the 1990s: the Framingham Heart Study. Circulation. 2006;113: 2914–2918.

12. Son C, Hosoda K, Matsuda J, et al. Up-regulation of uncoupling protein 3 gene expression by fatty acids and agonists for PPARs in L6 myotubes. Endocrinology. 2001; 142:4189–4194.

13. Ferré P. The biology of peroxisome proliferator–activated receptors: relationship with lipid metabolism and insulin sensitivity. Diabetes. 2004;53:S43–S50.

14. Tontonoz P, Spiegelman BM. Fat and beyond: the diverse biology of PPAR-gamma. Annu Rev Biochem 2008;77:289–312.

15. Na HK, Surh YJ. Peroxisome proliferator-activated receptor gamma (PPAR-gamma) ligands as bifunctional regulators of cell proliferation. Biochem Pharmacol 2003;66:1381–91.

16. Heikkinen S, Auwerx J, Argmann CA. PPARgamma in human and mouse physiology. Biochim Biophys Acta 2007;1771:999–1013.

17. Dussault I, Forman BM. Prostaglandins and fatty acids regulate transcriptional signaling via the peroxisome proliferator activated receptor nuclear receptors. Prostaglandins Other Lipid Mediat 2000;62:1–13.

18. Kliewer SA, Sundseth SS, Jones SA, Brown PJ, Wisely GB, Koble CS, et al. Fatty acids and eicosanoids regulate gene expression through direct interactions with peroxisome proliferatoractivated receptors alpha and gamma. Proc Natl Acad Sci USA 1997;94:4318–23.

19. Lehmann JM, Moore LB, Smith-Oliver TA, Wilkison WO, Willson TM, Kliewer SA. An antidiabetic thiazolidinedione is a high affinity ligand for peroxisome proliferator-activated receptor gamma (PPAR gamma). J Bio Chem 1995;270:12953–56.

20. Cho N, Momose Y. Peroxisome proliferator-activated receptor gamma agonists as insulin sensitizers: from the discovery to recent progress. Curr Top Med Chem 2008;8:1483–507.

21. Balakumar P, Kathuria S. Submaximal PPARgamma activation and endothelial dysfunction: new perspectives for the management of cardiovascular disorders. Brit J Pharmacol 2012;166:1981–92.

22. Schupp M, Clemenz M, Gineste R, Witt H, Janke J, Helleboid S, et al. Molecular characterization of new selective peroxisome proliferator-activated receptor gamma modulators with angiotensin receptor blocking activity. Diabetes 2005;54:3442–52.

23. Higgins LS, Depaoli AM. Selective peroxisome proliferator-activated receptor gamma (PPARgamma) modulation as a strategy for safer therapeutic PPAR-gamma activation. Am J Clin Nut 2010;91:72S–267S.

24. Beutler JA. Natural products as a foundation for drug discovery. Curr Protoc Pharmacol 2009. Chapter 9:Unit 9 11.

25. Cragg GM, Newman DJ. Natural products: a continuing source of novel drug leads.Biochim Biophys Acta 2013;1830:3670–95.

26. Henrich CJ, Beutler JA. Matching the power of high throughput screening to the chemical diversity of natural products. Nat Prod Rep 2013;30:1284–98.

27. Clardy J, Walsh C. Lessons from natural molecules. Nature 2004;432:829–37.

28. Ertl P, Roggo S, Schuffenhauer A. Natural product-likeness score and its application for prioritization of compound libraries. J Chem Inf Model 2008;48:68–74.

29. Sugano, M.; Koba, K.; Tsuji, E. Health benefits of rice bran oil. Anticancer Res. 1999, 19, 3651–3657.

30. Islam, M.S.; Murata, T.; Fujisawa, M.; Nagasaka, R.; Ushio, H.; Bari, A.M.; Hori, M.; Ozaki, H. Anti-inflammatory effects of phytosteryl ferulates in colitis induced by dextran sulphate sodium in mice. Br. J. Pharmacol. 2008, 154, 812–824.

31. Islam, M.S.; Nagasaka, R.; Ohara, K.; Hosoya, T.; Ozaki, H.; Ushio, H.; Hori, M. Biological abilities of rice bran-derived antioxidant phytochemicals for medical therapy. Curr. Top. Med. Chem. 2011, 11, 1847–1853.

32. Panlasigui, L.N.; Thompson, L.U. Blood glucose lowering effects of brown rice in normal and diabetic subjects. Int. J. Food Sci. Nutr. 2006, 57, 151–158.

33. Mohan, V.; Spiegelman, D.; Sudha, V.; Gayathri, R.; Hong, B.; Praseena, K.; Anjana, R.M.; Wedick, N.M.; Arumugam, K.; Malik, V.; et al. Effect of brown rice, white rice, and brown rice with legumes on blood glucose and insulin responses in overweight Asian Indians: A randomized controlled trial.Diabetes Technol. Ther. 2014, 16, 317–325.

34. Wang, O., Liu, J., Cheng, Q., Guo, X., Wang, Y., Zhao, L., et al., 2015. Effects of ferulic acid and γ -oryzanol on metabolic syndrome in rats. PloS One 10, 1–14.

35. Francisqueti, F.V., Minatel, I.O., Ferron, A.J.T., Bazan, S.G.Z., Silva, V. dos S., Garcia, J. L., et al., 2017a. Effect of gamma-oryzanol as therapeutic agent to prevent cardiorenal metabolic syndrome in animals submitted to high sugar-fat diet. Nutrients 9, 1–10.

36. Son, M.J., Rico, C.W., Nam, S.H., Kang, M.Y., 2011. Effect of oryzanol and ferulic acid on the glucose metabolism of mice fed with a high-fat diet. J. Food Sci. 76, 4–7.

37. H.M. Berman, J. Westbrook, Z. Feng, G. Gilliland, T.N. Bhat, H. Weissig, I.N. Shindyalov, P.E. Bourne, The Protein Data Bank (2000) Nucleic Acids Research 28: 235–242

38. Sheu SH, Kaya T, Waxman DJ, Vajda S. Exploring the binding site structure of the PPAR gamma ligand-binding domain by computational solvent mapping. Biochemistry. 2005 Feb 1;44(4):1193–209.

39. Kim S, Thiessen PA, Bolton EE, Chen J, Fu G, Gindulyte A, Han L, He J, He S, Shoemaker BA, Wang J, Yu B, Zhang J, Bryant SH. PubChem Substance and Compound databases. Nucleic Acids Res. 2016 Jan 4;44(D1):D1202–13.

40. The PyMOL Molecular Graphics System, Version 2.0 Schrödinger, LLC.

41. Morris, G. M., Huey, R., Lindstrom, W., Sanner, M. F., Belew, R. K., Goodsell, D. S. and Olson, A. J. (2009) Autodock4 and AutoDockTools4: automated docking with selective receptor flexiblity. J. Computational Chemistry 2009, 16: 2785-91.

42. Forli, S., Huey, R., Pique, M. E., Sanner, M. F., Goodsell, D. S., & Olson, A. J. (2016). Computational protein-ligand docking and virtual drug screening with the AutoDock suite. Nature Protocols, 11(5), 905–919.

43. J. Eberhardt, D. Santos-Martins, A. F. Tillack, and S. Forli. (2021). AutoDock Vina 1.2.0: New Docking Methods, Expanded Force Field, and Python Bindings. Journal of Chemical Information and Modeling.

44. O. Trott, A. J. Olson, AutoDock Vina: improving the speed and accuracy of docking with a new scoring function, efficient optimization and multithreading, Journal of Computational Chemistry 31 (2010) 455–461

45. Maier, J. A., Martinez, C., Kasavajhala, K., Wickstrom, L., Hauser, K. E., & Simmerling, C. (2015). ff14SB: Improving the Accuracy of Protein Side Chain and Backbone Parameters from ff99SB. Journal of Chemical Theory and Computation, 11(8), 3696–3713.

46. Sousa Da Silva, A. W., & Vranken, W. F. (2012). ACPYPE - AnteChamberPYthon Parser interfacE. BMC Research Notes

47. Bussi, G., Donadio, D., & Parrinello, M. (2007). Canonical sampling through velocity rescaling. Journal of Chemical Physics, 126(1), 014–101.

48. Parrinello, M., & Rahman, A. (1981). Polymorphic transitions in single crystals: A new molecular dynamics method. Journal of Applied Physics, 52(12), 7182–7190.

49. Schäfer, T., Kriege, N., Humbeck, L. et al. Scaffold Hunter: a comprehensive visual analytics framework for drug discovery. J Cheminform 9, 28 (2017).

